# Improving the Scalability of Bayesian Phylodynamic Inference through Efficient MCMC Proposals

**DOI:** 10.1101/2025.06.18.660471

**Authors:** Remco R. Bouckaert, Paula H. Weidemüller, Luis R. Esquivel Gomez, Nicola F. Müller

## Abstract

In Bayesian phylodynamics, we jointly reconstruct the posterior distribution of timed phylogenetic trees, evolutionary and population dynamic parameters. Most approaches in the field use Markov chain Monte Carlo (MCMC) for inference. Inferring timed phylogenetic trees using MCMC relies on a set of proposal distributions – operators – to explore different phylogenetic tree topologies, node ranks, and heights. However, inefficiencies in these proposal distributions limit the speed and scale with which phylodynamic analyses can be performed. Here, we introduce two classes of operators that allow scaling of phylodynamic analyses by reducing the time needed to reach convergence. We propose an improved proposal distribution to scale trees based on the intervals between consecutive nodes in the tree, and a set of operators informed by a pseudo parsimony score to explore different tree topologies. Contrary to already existing operators that make large changes to areas of the tree with no mutations at all, our new parsimony-focused operators target areas of the tree with many mutations.

We first demonstrate the correctness of these proposal distributions using a well-calibrated simulation study. We then demonstrate the improvements of using the updated proposal distributions using a number of large real-world datasets, showing a substantial increase in effective sample size per hour across viral datasets as well as a reduction in the number of samples required for the MCMC chain to reach stationarity. Finally, we apply the operators to a 10,000-sequence H3N2 dataset including phylogeography and finish the analysis within two weeks. The operators are implemented in the TargetedBeast package, which is available as an open-source package for BEAST2 and a user-friendly graphical user interface. The speed improvements in the TargetedBeast package are beneficial to any phylodynamics method implemented in BEAST2 that relies on standard phylogenetic trees, such as coalescent-skyline, birth-death skyline, and structured phylodynamic methods.

## Introduction

Bayesian phylogenetics has given us insight into the transmission dynamics of pathogens, the evolution of species, and even languages. Historically, data availability has been a major bottleneck. However, with the advent of next-generation sequencing, the inference algorithms of Bayesian phylodynamics are increasingly the main constraint to perform analyses. Most Bayesian phylodynamics inference software, such as MrBayes, RevBayes, or BEAST (7; 19; 35; 37), rely on Markov chain Monte Carlo (MCMC) to perform inference. MCMC is based on a random exploration of the state space, where a new state is proposed based on the current state through MCMC proposals, also known as operators, kernels, proposal distributions, etc. In line with the terminology of BEAST2, we will mainly use the term operator here. The phylogenetic state space grows superexponentially in size with the number of taxa, making the random exploration of tree space inefficient. As a result, using MCMC is limited to around a few thousand sequences unless tree space is restricted.

Attempts to scale up MCMC to larger datasets generally focus on two areas: speeding up the calculation of the posterior and improving the efficiency of the exploration of the state space. Calculating the tree likelihood (15) using Felsenstein’s pruning algorithm is often the most time-consuming part when calculating the posterior of a state, even when caching calculations. The BEAGLE library (3) aims to increase this calculation by exploiting hardware capabilities of CPUs and GPUs and is integrated with the most popular phylogenetic software packages. Delphy (39) takes another approach by explicitly representing all mutation events instead of integrating them out as in Felsenstein’s algorithm. This dramatically reduces the demand for computing the tree likelihood and allows fast computation of MCMC steps and analysis of large datasets. However, these speed improvements are currently limited to scenarios where there are few mutations, and the tree prior is simple and fast to calculate, such as constant or exponential coalescent. Its performance can deteriorate when there are a large number of mutations (e.g., when an outgroup is included in the analysis). Further, more complex tree priors, such as structured methods like the structured birth-death (21) or coalescent (27; 40), tend to require more complex calculations. As a result, the tree likelihood calculation is no longer the bottleneck, and speedups in likelihood calculations therefore will not translate to dramatic speedups overall.

Improving the exploration of state space includes parallel tempering (1; 25), a computationally intensive approach that runs several MCMC chains in parallel and periodically swaps states between chains. Bactrian proposals (38; 46) improve the efficiency of MCMC proposals that move a location or scale parameter over Gaussian proposals by recognising that the proposals near the mode of a Gaussian proposal (which is zero) do not help much in exploration of the state space. Bactrian proposals have two modes away from zero, and usually increase efficiency by at least 10% at no additional computational cost. However, this only provides a moderate improvement over non-Bactrian proposals. Joint operation on multiple parameters can additionally be used to improve convergence (5). Lastly, dataset-specific weighing of different operators can improve efficiency (10). Since the most suitable weighing cannot be established a priori before running the analysis, an adaptable operator was introduced that selects from among a set of proposals and during the MCMC records the acceptance probability of each proposal, the amount of change in the parameter of interest and the amount of calculation time to compute the posterior after a proposal. Based on these statistics, more efficient proposals are selected more often. Although it reliably selects more efficient proposals, it does not improve the efficiency of the best proposal in the mixture. Bouckaert (8) designed a set of operators specifically for moving groups of internal node heights in a tree, improving the efficiency of tree proposal in particular for non-ultrametric trees. All of these approaches increase the efficiency of MCMC proposals, but still do not allow us to scale up to thousands of taxa for a wide range of models.

Alternative inference methods to MCMC for inferring posterior distributions in Bayesian phylodynamics include nested sampling (36) and sequential Monte Carlo (41). However, they are often still outperformed by MCMC. Hamiltonian Monte Carlo (HMC) proposals leverage principles from Hamiltonian dynamics to propose new states in the Markov chain (4; 16). So far, HMC proposals for moving trees (34) suffer from larger computational overhead, making them not more efficient than the much simpler MCMC proposals. Variational approaches (9; 47) are perhaps the most promising for scaling up phylogenetic inference, but only offer approximations to the posterior and have so far not been proven to scale well.

In order to deal with large datasets in practice, these datasets are subsampled, resulting in a loss of resolution of the features of interest, and/or pipelines are crafted to combine individual analyses of subsets (see, for example, (12)), or by splitting datasets into multiple smaller trees and then perform phylodynamic inference jointly from all of them (29).

Here, we provide a number of improvements to the MCMC operators specifically targeted at trees informed by the number of mutations in regions of the tree. They aim to more efficiently move all node heights in a tree, as well as target specific regions in a tree where topology changes are expected to lead to larger average jumps in tree space. We take the widely used mixture of BEAST2 operators as baseline, which already incorporates best practice techniques like Bactrian proposals (46; 38), adaptive operator sampling (10), and advanced tree stretching (8). Together, our new set of operators dramatically improves the speed at which we can reconstruct pathogen outbreaks over this baseline, particularly for large datasets. The proposals are implemented in BEAST2 (7) in the open source TargetedBeast package available from https://github.com/nicfel/targetedbeast/ licensed under the LGPL 2.1.

## Methods and Materials

### Weighted tree proposals

Previous work has shown that the convergence of Bayesian phylogenetic inference can be improved by using parsimony-guided tree proposals (48). Here we build upon this idea by using weighted and targeted tree proposals. To do so, we first compute what we call the pseudo-parsimony sequence of every internal node, using all parsimony-informative sites in the data. At each node, we compute a consensus sequence of the nodes below using an upwards pass (see Figure 1A). At each position, a node can be either A, T, G or C, any pairwise combination of the two, or N.

**Figure 1.**
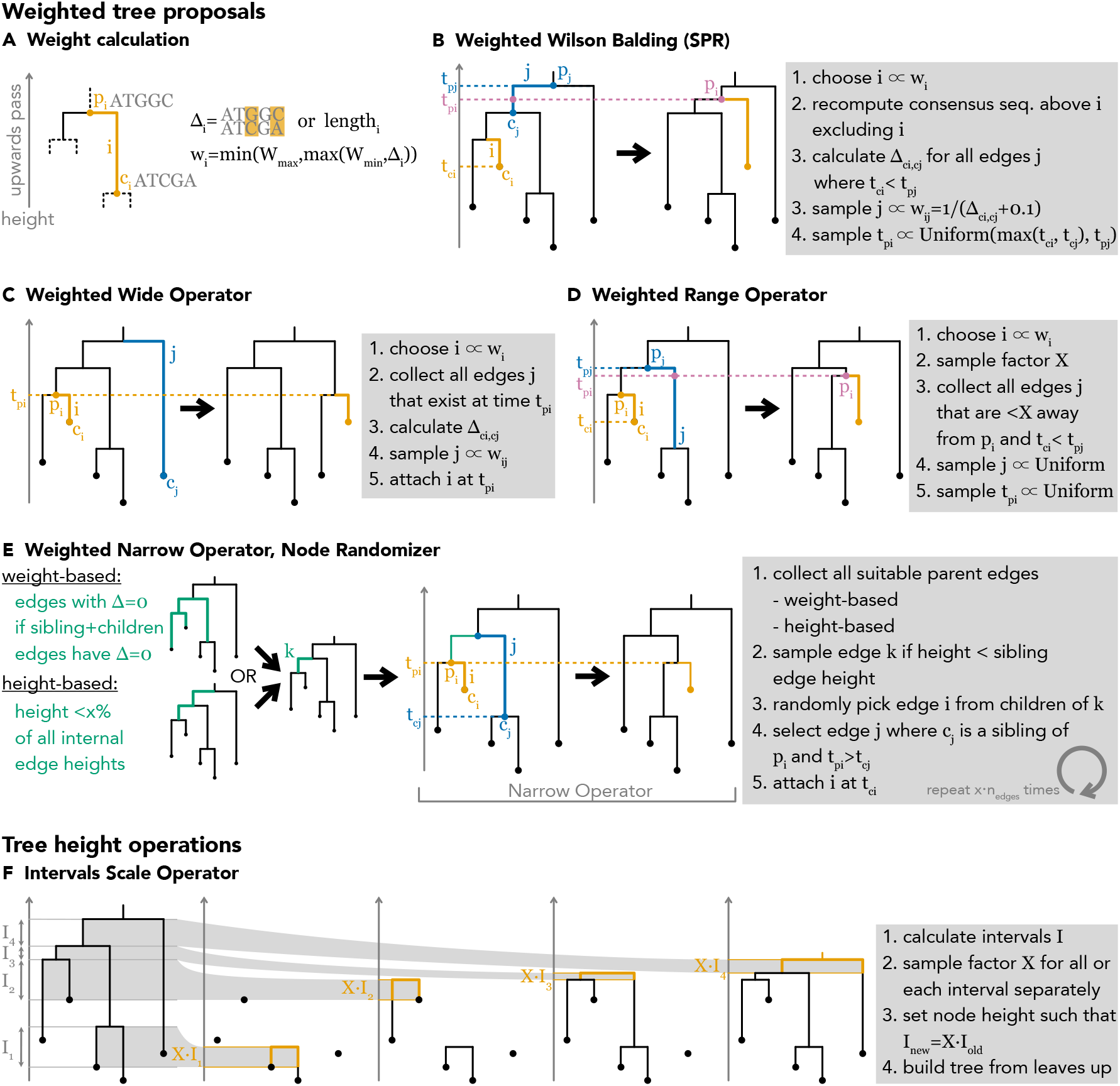
Schematic descriptions of the operators used in the TargetedBeast package. Here, we show a schematic representation of the pseudo-parsimony calculation used in the weighted tree operations and the principle of the interval scale operator. *i* generally refers to the edge that is operated on, while *j* generally refers to the edge where *i* is reattached to. *p*_*i*_ refers to the parent node of edge *i*, while *c*_*i*_ refers to the child node of edge *i*.

For each edge *i*, we compute the difference between the parent-child consensus nodes Δ_*i*_. The weight of edge *i*, w_*i*_, to be operated on is computed using:

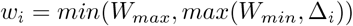

The minimum weight (*W*_*min*_) is used such that edges with Δ_*i*_ = 0 are still operated on and such that moves leading to Δ_*i*_ = 0 are not rejected by default. The maximum weight (*W*_*max*_) is to avoid operators operating almost exclusively on edges with large Δ_*i*_, i.e., long edges in the tree.

This enables us to approximate the number of mutations on each branch and to compute distances between nodes as the number of consensus positions that differ between them. This differs from a full parsimony reconstruction in that there is only an upwards (from tips to root), but not a downwards pass (from root to tips) involved in computing the consensus sequences. This is done to reduce the requirements for updating the consensus nodes when updating the tree topology. As the consensus sequences are only used to propose new tree topologies and not directly in the posterior calculation, this approximation does not affect the inference results. The ‘edge weights’ class is implemented such that it can be easily extended using different ways to weight which edges to operate on and where to compute distances between them. The edge weights are then used in a series of tree operators similar to existing tree operators implemented in BEAST and BEAST2.

#### Weighted Wilson Balding

This operator is an adaptation of the subtree-prune-regraft (SPR) move (43). First, an edge to operate on is randomly chosen with probability proportional to its weight *w*_*i*_. Then, the consensus sequences of any nodes above edge *i* are recomputed without edge *i*. Second, the distance of the consensus sequences between *c*_*i*_ and any node *c*_*j*_, Δ_*ij*_, for every edge *j* for which the height of *p*_*j*_, the parent node of *j*, is larger than *c*_*i*_ is computed. Third, we use 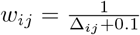 as the weights to sample which edge *j* to reattach edge *i* to. 0.1 is added such that the denominator never becomes 0. Fourth, the new height of *p*_*i*_ on edge *j* is sampled randomly from a uniform distribution between *p*_*j*_ and the older of *c*_*i*_ and *c*_*j*_ (see Figure 1B). Additionally, we have an alternative formulation of this operator where we use the length of each edge as the edge weights *w*_*i*_. This is to preferentially select edges to operate on that tend to contribute disproportionately to the tree length and tree prior instead of the tree likelihood.

#### Weighted Wide Operator

First, an edge to operate on is randomly chosen with probability proportional to its weight *w*_*i*_. Then, we collect a list of all edges *j* that are coexisting with the height of the parent of edge *i, p*_*i*_. For that list of edge *j*, we then compute the distance of the child of *j* to the child of *i* Δ_*ij*_. We then use 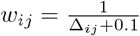 as the weights to sample which edge *j* to reattach edge *i* to. The reattachment time on *j* is equal to *p*_*i*_ (see Figure 1C). This is largely the same as the SPR move described above, but does not change the tree length and reduces the need to compute distances between consensus sequences.

#### Weighted Range Operator

First, an edge to operate on is randomly chosen with probability proportional to its weight *w*_*i*_. Then, we randomly sample a value X from a Bactrian distribution, i.e., a mixture of two Normal distributions that are symmetrically displaced from 0. We then collect all edges *j* that are X away from *p*_*i*_, irrespective of which direction it is from *p*_*i*_ (see Figure 1D). This has been described as the “Sub-tree Leap Operator” (14) in https://github.com/beast-dev/beast-mcmc/blob/master/src/dr/evomodel/operators/SubtreeLeapOperator.java. We used the subtree leap approach over the subtree slide, and removed the narrow operators from the XML setup. We also tested a version of the subtree leap operator that uses the distances to any potential target edge *j* as described for the SPR move to select target edges from a weighted distribution. However, the additional computations needed did not seem to lead to an improvement in convergence.

#### Node Randomizer

Each narrow exchange move re-attached and *i* to the edge *j* if the grandparent of *i* is equal to the parent of *j*. Instead of a single narrow operation, we use a series of narrow operations to make larger jumps in tree space (see Figure 1E). The operator comes in two forms. In the first one, the **height-based** version, we perform an operation on *x*% of all edges in the tree, with *x* being a tunable/adaptable parameter. We first sort all edges by length, and then choose the length of the *x*^*th*^ percentile as the threshold for all operations. Among all edges *k* that have an edge length below the *x*^*th*^ percentile edge length, we perform *x* times the total number of edges narrow operations.

In the **weight-based** version, we perform narrow operations on all edges for which Δ_*i*_ = 0 for all edges involved in the move. We perform this operation *x* times the total number of edges, with *x* being a tunable parameter, as for the height-based version.

### Intervals scale operators

Traditionally, BEAST changed the heights of all nodes by scaling them with a scale factor drawn from some distribution. However, this can lead to poor performance when working with serially sampled phylogenies, as scaled node heights can be inconsistent with sampling times, leading to negative branch lengths being proposed leading to immediate rejection of the proposal. Tree stretch operators (8) avoid this issue and lead to improvements in convergence, though here we introduce an operator that is much more efficient.

To scale timed phylogenies, we follow the following process. First, we define a set of intervals as the height between a parent node and the maximum height of either child node. We then choose a scaler *X* randomly from the Bactrian distribution (46; 38). We next scale the length of each interval by *X*. Lastly, we rebuild the tree from the leaves up, setting the height of each node such that the new interval is *X* times the old interval (see Figure 1F). This approach is agnostic to whether child 1 or child 2 is the more ancestral node, and which child is more ancestral can change during a move. Related operators that scale the subtree below nodes have been implemented previously, such as in (20). This way of operating on the tree allows for the rank of nodes to change through a move, but keeps the topology the same. As all intervals will always have a length greater than or equal to 0, this move never proposes invalid trees, in contrast to the usual tree scale operator. To allow for simultaneous moves on the clock rate and the population size (or birth-death, and sampling rates in the case of the birth-death model), we keep track of the tree length before and after the move. We then simultaneously scale the clock rate and/or population size by the change in tree length.

We additionally implemented the same operator, where we chose *X* randomly for each interval. The interval scale operator is conceptually simple to extend to jointly operate on local clock rates, epoch clock models, birth-death-sampling rates, coalescent skyline parameters, and the edge lengths of timed, serially sampled phylogenetic trees.

### Viral Datasets

To test the performance of our newly implemented operators, we compiled a series of human viral datasets. We chose the datasets to represent a broad range of outbreak dynamics, sampling time intervals, sampling densities, and data availability. We used two long-term influenza datasets (influenza A/H3N2, HA, and influenza B, PB1) as described in (28), which were compiled from fludb. Both datasets were sampled over multiple decades. The influenza B dataset contained both Victoria and Yamagata lineages We compiled an additional dataset using influenza A/H3N2 sequences sampled between 2022-05-01 and 2025-01-28 from gisaid (13) (https://doi.org/10.55876/gis8.250604ec). Additionally, we compiled two datasets from Pathoplexus: an mpox viral dataset collected in the USA over 3 years (30) and a West Nile dataset (31) collected over several years. We subsampled the data using the scripts https://github.com/nicfel/TargetedBeast-Material/blob/main/Applications/convertPathoplexusMPOX.R and https://github.com/nicfel/TargetedBeast-Material/blob/main/Applications/convertPathoplexusWestNile.R.

## Data availability

The source code for the analyses performed, such as the R scripts to recreate figures, is available here https://github.com/nicfel/targetedbeast-material. R-based analyses were conducted using `ggplot2` (42), `gridExtra` (2), `colorblindr` (23), and `coda` (32) packages for data manipulation, visualization, and MCMC convergence diagnostics. The new operators are implemented as part of a BEAST2 package called TargetedBeast. The source code for this package is available here https://github.com/nicfel/targetedbeast.

## Results

### Well-calibrated simulation study

To ensure a level of confidence in the correctness of the implementation, we conducted a well-calibrated simulation study (24). First, we sampled 100 trees with *n* taxa using MCMC with tip dates fixed at randomly selected times uniformly drawn from the unit interval. A coalescent with constant population size was used as a tree distribution where the population size was drawn from a narrow log normal distribution (*S* = 0.05) and mean such that the tree height remained small on average (2.90 for *n* = 50, 2.85 for *n* = 100, 2.01 for *n* = 500).

Next, we simulated an alignment on the tree with *k* sites under an HKY model with *κ* drawn from a lognormal (*M* = 1, *S* = 1.25) and frequencies from a Dirichlet (*α* = 4, 4, 4, 4) using gamma rate heterogeneity (44) with shape exponentially distributed (*λ* = 1). A strict clock with a clock rate of 1.0 was used. Then, we ran analyses for each of the 100 trees in BEAST2, where the tree, as well as the population size, were estimated. We measure how often a true parameter value is in the 95% highest probability density (HPD) interval, which for 100 runs should be in the range 91 to 99 to be acceptable.

We ran the analysis with *n* = 50, 100 taxa and a selection of *k* = 25, 50, 100, 500 sites to get a wide range of data sets. Coverage of the various parameters in the analysis is shown in Table 1, which should be in the range 91 to 99 for 95% of the entries. Since just two out of 80 entries are outside this range, it suggests that the coverage is sufficient and does not give a reason to believe that the operators have been incorrectly implemented.

**Table 1:** Coverage of parameters (shown in the first column) in the well-calibrated simulation study. Coverage is defined as the number of runs where the true value is included in the 95% HPD interval and is expected to be in the range of 91 to 99. Any values that fall outside this range are bolded.

| Number of sites | 50 taxa |  |  |  | 100 taxa |  |  |  |
| --- | --- | --- | --- | --- | --- | --- | --- | --- |
|  | 25 | 50 | 100 | 500 | 25 | 50 | 100 | 500 |
| tree height | 91 | 96 | 92 | 97 | 98 | 94 | 93 | 95 |
| tree length | 94 | 97 | 95 | 95 | 98 | 98 | 91 | 93 |
| kappa | 96 | 91 | 96 | 99 | 98 | 96 | 96 | 95 |
| frequencies A | 95 | 95 | 91 | 94 | 95 | 95 | 95 | 98 |
| frequencies C | 94 | 93 | 93 | **90** | 92 | 93 | 97 | 94 |
| frequencies G | 93 | 97 | 94 | 92 | 94 | 94 | 93 | 92 |
| frequencies T | 94 | 96 | 95 | 96 | 97 | 94 | 92 | 95 |
| gamma | 98 | 94 | **100** | 95 | 95 | 97 | 93 | 94 |
| pop | 96 | 97 | 96 | 96 | 98 | 94 | 94 | 96 |
| coalescent | 95 | 94 | 95 | 91 | 93 | 94 | 91 | 92 |

Furthermore, since the new operators affect topology, we checked that the posterior clade support estimates averaged over all runs coincide with the true clades (24). To this end, we divided the clades into 10 bins, where the first bin contains all clades with less than 10% support, the second bin 10-20% support, etc. Then, for each bin, we checked how many of these clades are in the tree from the ground truth. If clade support estimates are correct, a bin with *x, x* + 10% support should have a fraction *x* + 5% of the clades in the ground truth trees. Figure S2 shows support for the trees with 100 taxa and 500 sites, but other configurations look very similar. This further supports that operators have been correctly implemented. We further performed a validation of the implementation of TargetedBeast in Text S1 using a real dataset of influenza B sequences.

### Improved convergence analysing real-world, viral datasets

One of the measures used most frequently to assess convergence of MCMC chains is the effective sample size (ESS). Two of the most crucial quantities to determine how rapidly a chain converges are the ESS per unit of time and how rapidly the stationary phase of an MCMC chain is reached. The latter quantity is sometimes referred to as the burn-in phase of an MCMC chain. To verify that the new operators contribute to the improvement of ESS per hour, we considered five real-world sequence alignments sampled over different time periods. We used two datasets of the HA segment of Influenza A/H3N2. The long-term dataset was sampled over multiple decades. The short-term dataset contained sequences sampled over 3 years in North America. The influenza B dataset contains sequences from the PB1 segments sampled over multiple decades. Influenza B underwent a diversification into Yamagata and Victoria lineages, leading to a very different tree shape than, for example, influenza A/H3N2. The MPox Virus dataset contained MPXV samples collected in the USA over multiple years. MPox is a double-stranded DNA virus with a genome of about 200kb. We further used a West Nile virus dataset sampled in the USA. All datasets consisted of 1000 sequences, except the MPXV dataset, which, after multiple filtering steps of the available data, contained 733 isolates. We chose these five human viral datasets to test the performance of the new operators under a range of scenarios, such as different genome sizes, rates of evolution, and different sampling intervals, all combining to very different tree shapes, as highlighted by Figure S5. Each dataset was run using a strict clock model and an *HKY* + Γ_4_ site model.

We tested three scenarios: default settings with site model parameters reweighted, a scenario where interval operators replace tree stretch and epoch flex operators, and a scenario where all new operators are included. Each run was started using the UPGMA starting tree in BEAST2, and not a random starting tree, as is the default setting. Figure S3 shows the change in ESS per hour for posterior, likelihood, and prior between these three scenarios. Each dataset was run for 100 million iterations, with three independent chains. Note that the y-axis has a log scale.

We next compared the time it took for the MCMC chain to reach the stationary phase. Here, we define reaching the stationary phase as the first 10 logged iterations that are within the 95% confidence interval of the converged chain. As shown in Figure S4, we find large reductions in the burn-in time required for the MCMC chains.

To provide a tangible estimate of the overall improvement between reduction in burn-in and faster convergence, we compare how long it took each chain to reach an ESS of the posterior of at least 100 and find large reductions in the time needed across all datasets (Figure 2).

**Figure 2.**
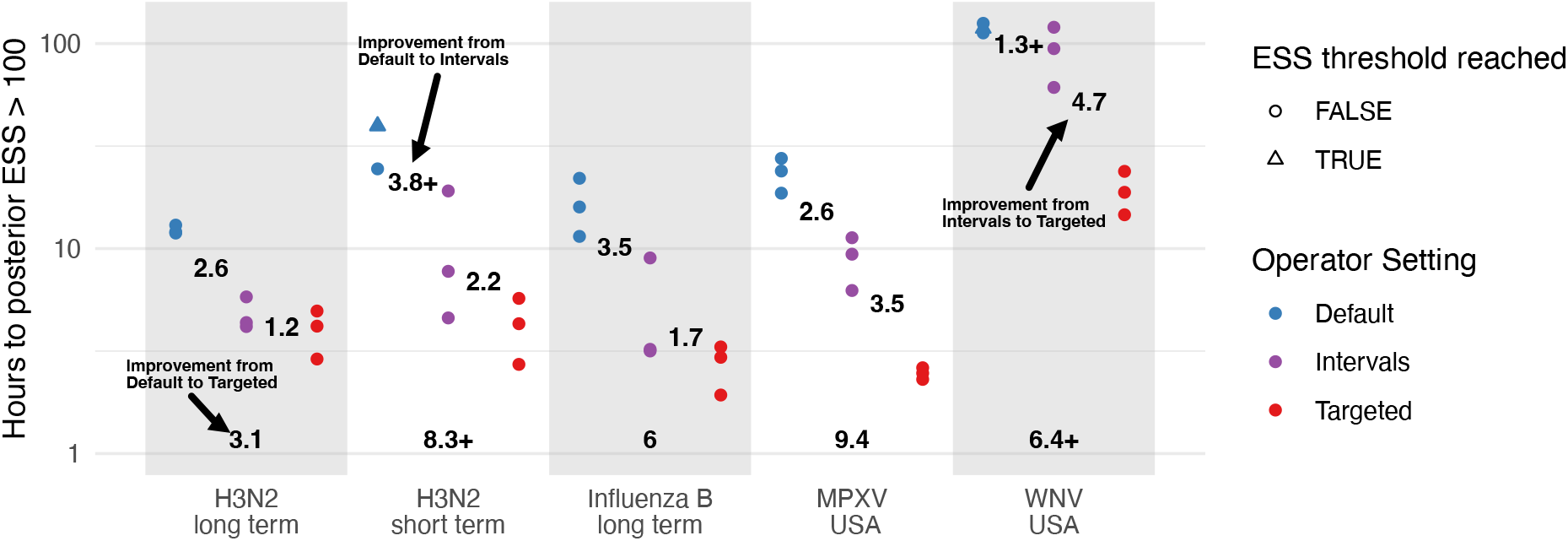
Time it took individual runs to reach a posterior ESS of at least 100. For each dataset on the x-axis and operator setup highlighted by the different colors, we show how long it took each run to reach a posterior ESS of at least 100. The same is shown for the likelihood and prior in the supplement. The individual values denote the fold reduction in time to ESS *>* 100 between the different operator settings. For runs marked by a triangle, the ESS threshold was not reached during the 100 million iterations, and the reduction in time to ESS *>* 100 is an underestimate, highlighted by the + sign.

### Bayesian phylogeographic analyses of a 10,000 sequence influenza A/H3N2 dataset

To provide a tangible example of how this BEAST2 package enables larger analyses, we performed an analysis on 10,000 H3N2 influenza sequences from North America from 2023-2025 using a HKY+Γ_4_ substitution model, a strict clock model, and a constant coalescent tree prior. Further, we jointly performed a discrete trait (22) reconstruction using 7 different locations in North America. This analysis was not meant to capture all the complexities of the geographic spread of Influenza A/H3N2, but rather highlight some of the added capabilities of this approach and new types and scales of analyses that can be performed.

We ran 8 different chains on single cores for 10 days and then combined the chains to a total of 250 million iterations using BEAGLE (3) to accelerate tree likelihood calculations and using the TargetedBeast package for BEAST2 (7). All relevant ESS values were above 200, so we consider the analysis converged. In Figure 3, we show the maximum clade credibility tree of the 10,000 isolates analyses with the traces for the posterior, likelihood, and prior shown in Figure S6. This analysis highlights the type of analyses that are now feasible, but impractical without TargetedBeast.

**Figure 3.**
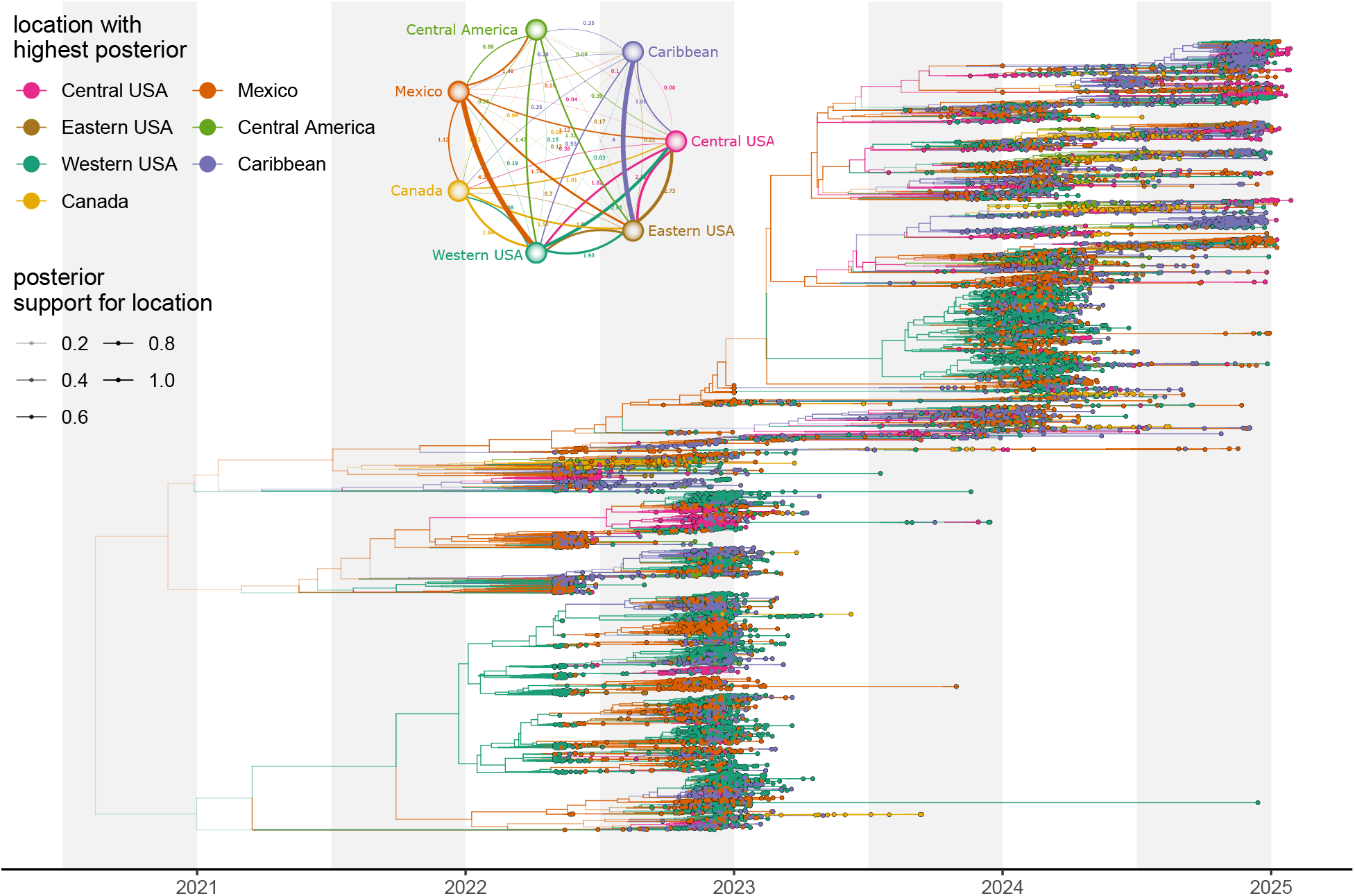
Joint Bayesian demographic and discrete trait analysis with 10,000 sequences of H3N2. Conditional clade credibility (CCD0) summary tree coloured by location with greatest posterior support. The color is shaded by posterior support value. Nodes with lower support are greyed out. Migration rate estimates jointly inferred with tree displayed as well.

We next estimated how the larger datasets contribute to a decrease in the uncertainty of the parameter estimates of the discrete phylogeographic model. To do so, we subsampled the 10,000 isolates dataset to 1000 isolates. We then compare the reduction in the ratio of upper over lower limit of the 95% HPD interval between the 1000-isolate dataset and the 10,000 isolate dataset. For the trait clock rate, the average rate of movement between regions, we find that he 95% HPD interval is reduced by about 50% (Figure S7).

## Discussion

We demonstrated that the new operators perform well on large datasets and improve mixing by anywhere from 3-fold to 10-fold. Furthermore, we performed a 10,000 sequence analysis in 10 days, which included discrete phylogeography (22) and is one of the largest fully Bayesian phylodynamic analyses we are aware of performed to date. The new operators are available for use with any of the phylogenetic models included in BEAST2 and are not limited to any particular models. The only assumption is that the analysis contains a single rooted time tree. This makes these new operators ideal for performing large-scale viral or bacterial analyses using coalescent, birth-death, and structured methods.

The topology-changing operators are, however, not always beneficial. These operators are informed by a pseudo-parsimony calculation to identify areas of interest in the tree, which can incur a relatively large overhead. Consequently, ESS per hour can suffer from degradation compared to when using the default operators. However, since small datasets usually run in a short period of time (in the order of hours, not days), such degradation may not be particularly problematic. In general, we expect larger increases for larger datasets. However, as running datasets with several thousand sequences is challenging to impossible using the default operator settings, the improved operator performance for these datasets are challenging to quantify.

In contrast, the interval operator never results in decreased performance and, in fact, often improves performance on small datasets. The overhead of the intervals operator is equal to that of the operators it replaces, but the proposals move all internal nodes in a tree more efficiently than other operators. The interval operators are readily extendable to jointly operate on time-varying coalescent or birth-death rates and the tree, using different scales for different parts of the tree. Likewise, these operators can be extended to include rates of relaxed clocks in such a way that evolutionary distances are not changed, which should lead to better mixing.

Apart from getting much better ESS per hour, the new operators also show a tendency to reduce burn-in considerably. Reduced burn-in is not only useful in standard MCMC analyses, but can also help speed up the variational approach as described in (9) and online analysis (6), which both rely on rapidly traversing burn-in.

We implemented the code such that it is readily extendable to different ways of weighting targets. Instead of parsimony-informed operators, it is possible to use pairwise sequence distances as follows. First, calculate a pairwise distance matrix and perform principal component analysis (PCA) or another dimension-reducing technique like multidimensional scaling to project all sequences to points in a *k*-dimensional space. Internal nodes in a tree are placed in the same *k*-dimensional space by recursively putting them on a line between the two child nodes, taking branch lengths into account. Weights *w*_*i*_ for node *i* are calculated as the distance to its parent node in the projected *k*-dimensional space. Likewise, for the Wilson Balding and wide exchange operators, the candidate positions *j* to place a clade are weighted 1*/*(0.1 + *d*_*ij*_) where *d*_*ij*_ is the distance from the clade node *i* to the candidate clade *j*.

Initial exploration of this PCA-based weighting scheme showed us that MCMC performance is comparable to the mutation-informed weighting scheme but performs better compared to pseudo parsimony on small data sets. The PCA-based scheme has less overhead since it does not require tracking all mutations in a phylogeny. So, while information is lost when using pairwise distances, the faster computation appears to compensate for this, resulting in comparable ESS per hour. Since there are a large number of implementation details, such as choice of distance metric, details of the dimension reducing technique, whether to take branch lengths and branch rates into account, setting of the minimum weight, etc., there is potential to get even better performance. Exploring this space of possibilities is outside the scope of this paper.

For both small and large datasets, operator weights (that is, the proportion of time the operator is selected to do a proposal in an MCMC chain) may need to be adjusted for the dataset at hand. These weights need to become stable throughout an MCMC run for the chain to reach stationarity. Adaptable operator samplers (10) go some way to learn optimal weights based on the amount of parameter change per unit of time for real-valued parameters, so they can assign weights among, for example, tree scalers and interval operators. However, from our experience, for topology changing operators, a metric like the Robinson-Foulds distance (33) appears to be too crude to distinguish between the various topology changing operator sets. Experimentation with more sophisticated metrics on tree space, like the ranked nearest neighbor interchange distance, may be beneficial in determining optimal proposal weights.

Instead of using the rudimentary pseudo-parsimony approach to calculate edge weights employed here, the implementation is readily extensible to explore different ways to calculate edge weights and distances. In particular, one could use a base tree, such as from a full parsimony or maximum likelihood inference, as a ‘guide-tree’ to perform operations, while still preserving full Bayesian inference.

Another area for exploration is for models not based on a single rooted time tree, such as the multi-species coalescent (17; 45) and network models (28; 26), where local trees are restricted to be included inside a species tree or network.

Overall, we provide a series of improvements to the current MCMC algorithm in BEAST2 that are straight-forward to utilize with most existing tools implemented in BEAST2 and a path forward to explore further efficiency improvements to enhance the scalability of Bayesian phylodynamics.

## Acknowledgements

We gratefully acknowledge all data contributors, i.e., the Authors and their Originating laboratories responsible for obtaining the specimens, and their Submitting laboratories for generating the genetic sequence and metadata and sharing via the GISAID Initiative, on which this research is based. We would like to thank Jiansi Gao for helpful discussions on the interval scale operator. NFM is funded in part by a UC Noyce Initiative Award and in part by the CDC CFA C-Core award. PHW is funded by a UC Noyce Initiative Award.

## Supplementary material

### Text S1: Detailed analysis of Influenza B using different operator weights

As a further comparison of the performance and to further validate the implementation of TargetedBeast, we performed an extensive analysis using the Influenza B dataset. To do so, we subsampled 198 sequences from the larger influenza B dataset described in *Viral Datasets*. To investigate the effect of the new set of operators, we set up analysis in BEAUti with HKY substitution model (*κ* ∼ log-normal(*m* = 1, *s* = 1.25)), with estimated frequencies (*π* ∼ Dirichlet(*α* = [4, 4, 4, 4])) and gamma rate heterogeneity (*γ* ∼ Exponential(*m* = 1)). A coalescent with constant population size (pop size ∼ 1*/X*) is used as a tree prior and a strict clock (clock rate~Uniform[0, ∞]) for the branch rate model. All priors are defaults in BEAST2, and operator weights were initially kept at default values as well. However, since this resulted in an unacceptable low effective sample size (ESS) for site model parameters compared to other parameters, weights were doubled for these operators so ESSs became acceptable.

This gives us three scenarios: 1) all default settings, 2) reweighted operators and 3) new operators (with site models operator weights the same as scenario 2). Since it is well known that ESS can vary between different MCMC runs, even with the same model (18), we ran the analysis with these three scenarios 50 times. Figure S1 shows the distribution of ESSs for these runs for various parameters. As a general rule of thumb, an ESS of 200 is widely used as a stopping criterion for phylogenetic MCMC analyses (11). ESS per unit of time is better when using our new operators for all items of interest except for *κ* and *γ* where the ESS is over 2000, so more than sufficient. The tree height and length ESSs improve massively with the new operators over those of the first two scenarios.

Since we ran long runs, as an extra check, we compared estimates of the default reweighted combined runs with the targeted runs. Results are shown in Table S1 and estimates are all consistent.

**Table S1:**
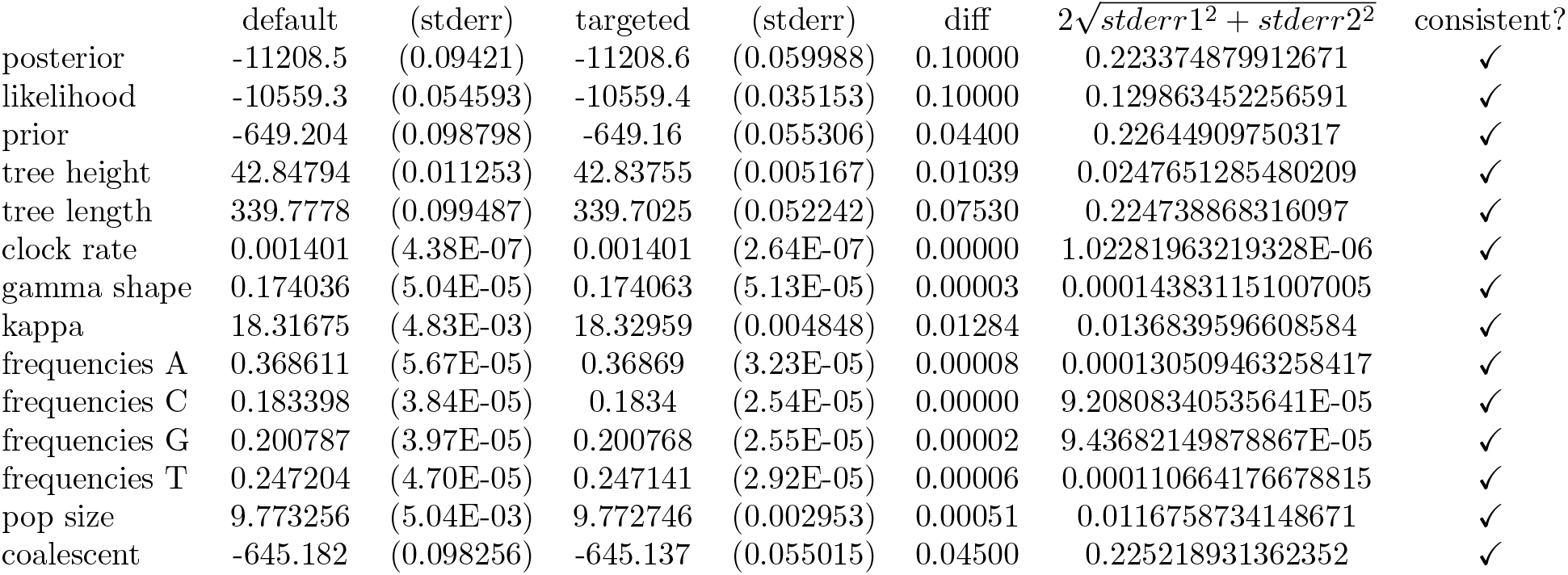
Estimates for parameters of the 50 runs combined for reweighted default operators and targeted operators obtained by the LogAnalyser tool from BEAST2. Numbers in brackets are the standard error for the estimates (note that for targeted operators, these are generally a bit smaller due to larger ESSs). The last column indicates whether the absolute difference between estimates is less than twice the square root of the sum of squared standard errors, i.e., whether |*default*−*targeted*| is less than 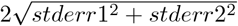. Since this is true for all estimates, all estimates can be considered to come from the same distribution, so both sets of operators are unbiased.

**Figure S1:**
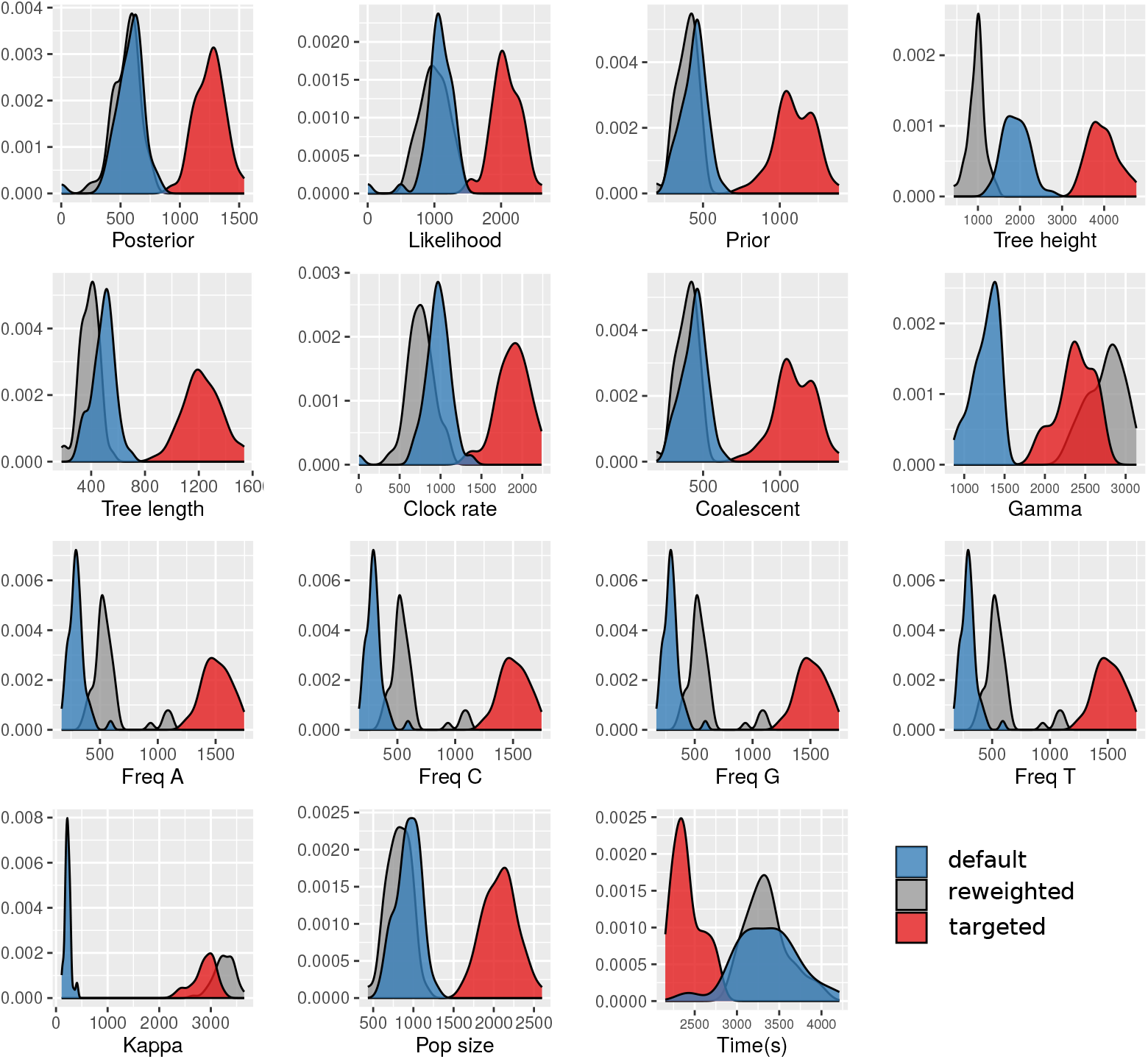
ESS distribution for parameters and different runs. All panels show the distribution of ESS for the various parameters involved in the analysis, with the x-axis showing the ESS and the y-axis the density, with the exception of the last panel, which shows run time in seconds. Blue curves represent default settings, grey curves are for reweighted operators, and red curves are for our new operators. For all cases, the new operators are never worse (unless exceeding 2000), often substantially better, but ESS per unit of time is always (much) better.

**Figure S2:**
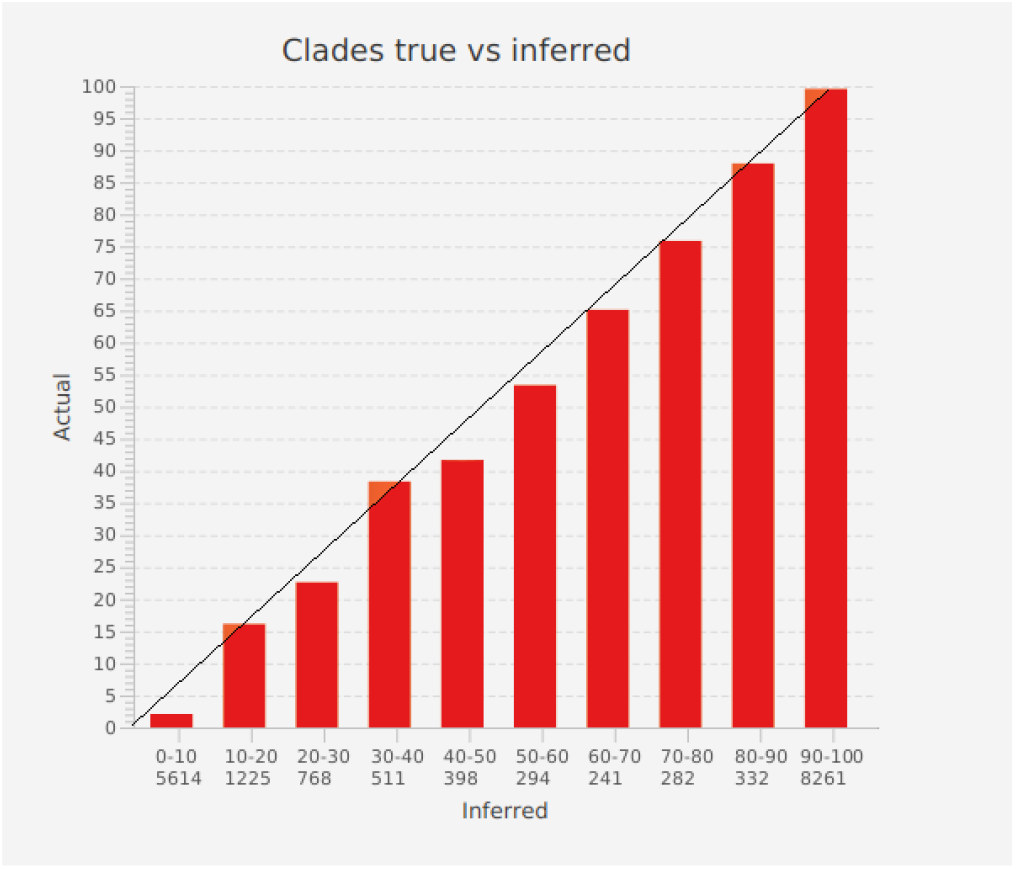
Clade support comparison. A bar graph is constructed by counting how many inferred probabilities fit in a bin, then calculating how many of the true values are in a certain bin. Each bar represents a 10% sized interval, where the first bar represents predictions in the range 0 to 10%, the second bar ranges from 10% to 20%, etc. The number below the range represents the number of predictions that fit in that range for each of the 100 posterior distributions. For example, there are 8261 predictions in the range 90-100% (last column) in the bar chart. The size of the bar is the percentage of times the true prediction is in the bin with the associated prediction. Bars are close to the diagonal x=y line, suggesting clade probability estimates are correct.

**Figure S3:**
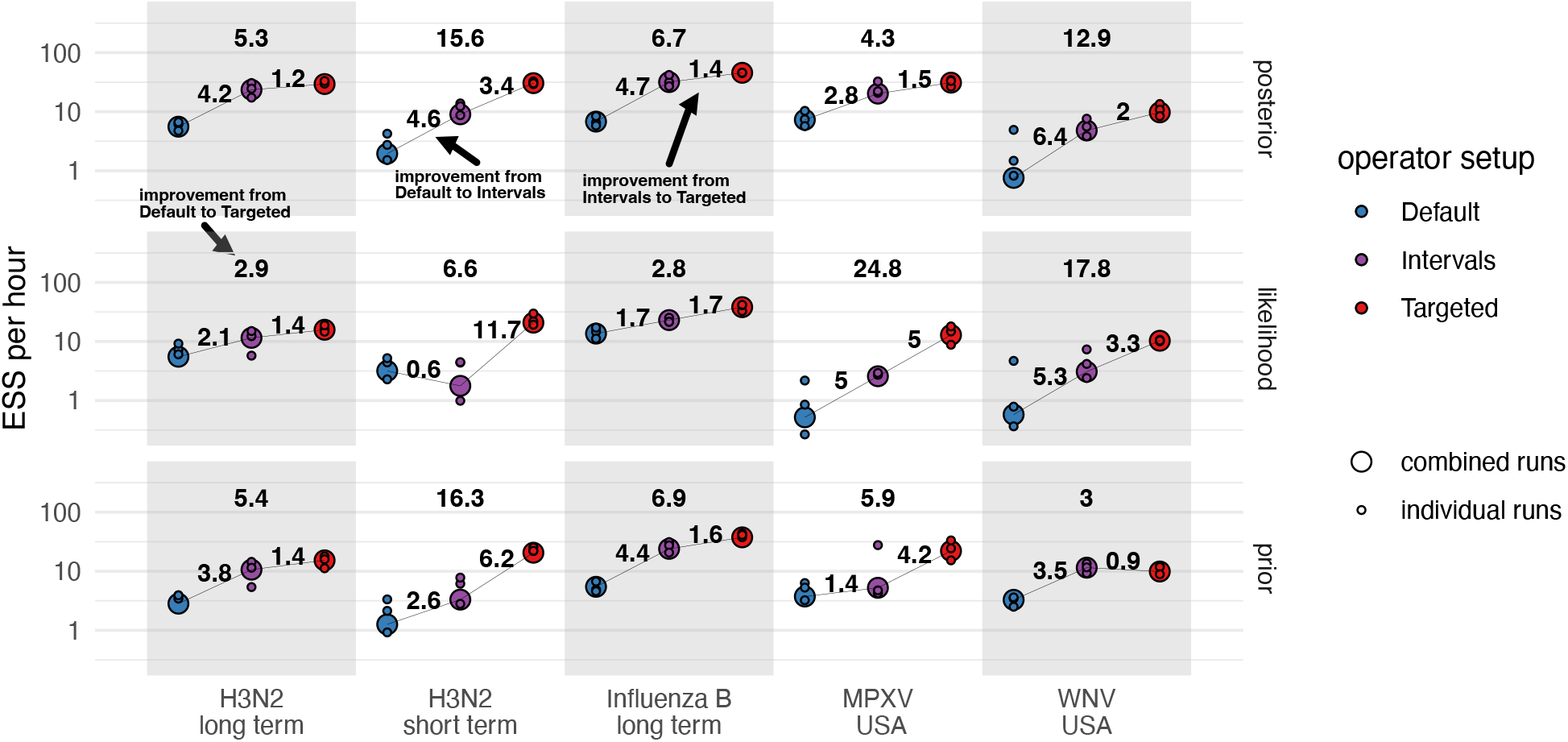
ESS per hour for posterior, likelihood, and prior values using different operator settings. For each dataset on the x-axis and operator setup highlighted by the different colors, we show the ESS per hour on a log scale on the y-axis. The ESS is shown for each of the three individual runs, as well as for the combined chain (large circles). The numbers show the fold improvements in ESS per hour when using the new operators. The individual fold increases are highlighted in the figure and made between Default and Targeted, Default and Intervals, and Intervals to Targeted. Each run was started using the UPGMA starting tree in BEAST2.

**Figure S4:**
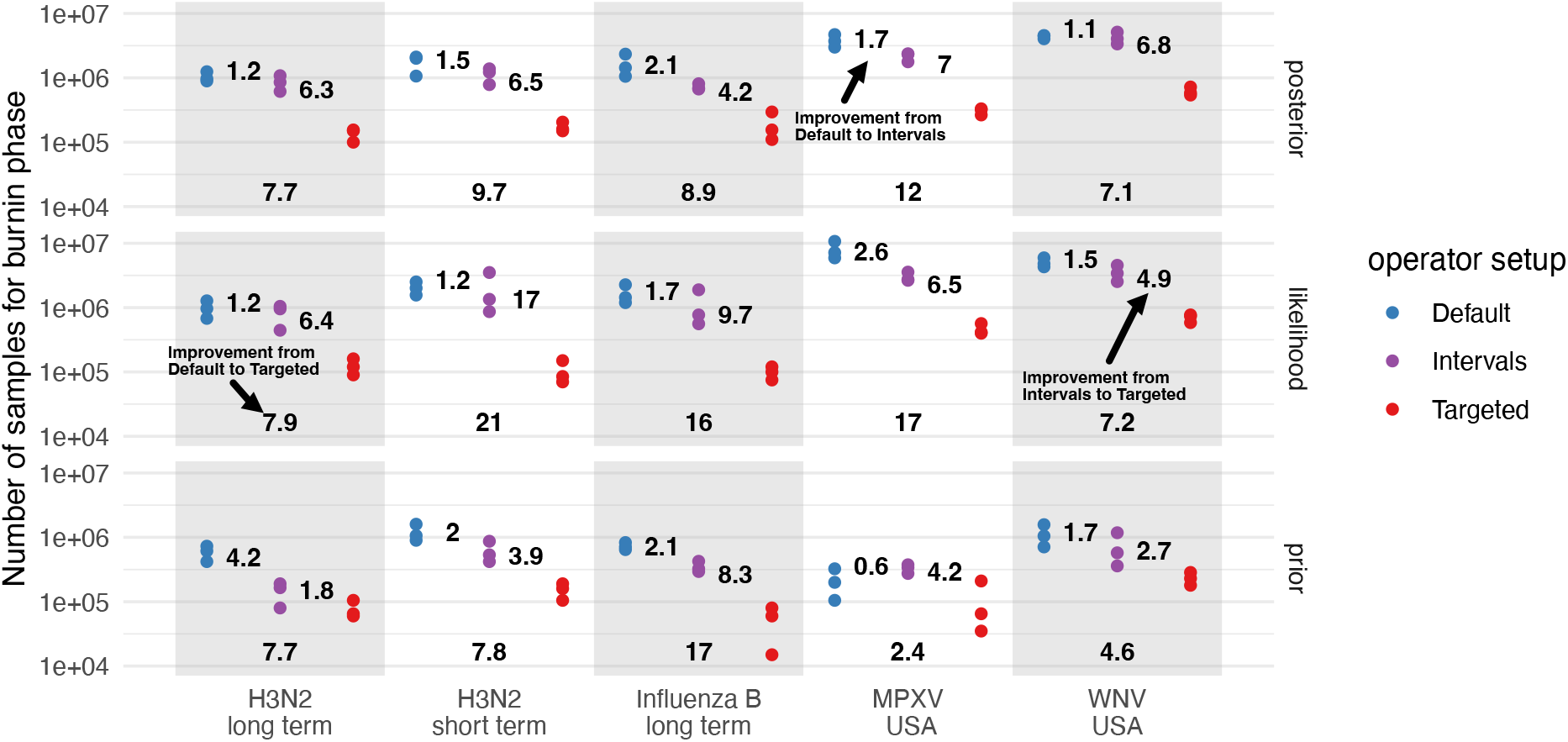
Reductions in the number of MCMC steps required until the stationary phase is reached. Here, we show the number of MCMC samples or steps on the y-axis required until the stationary phase is reached for the posterior probability value, the likelihood, and the prior probability for the different real world datasets. We here define the burn-in phase as the first ten consecutive logged iterations that are within the 95% HPD of the stationary phase.

**Figure S5:**
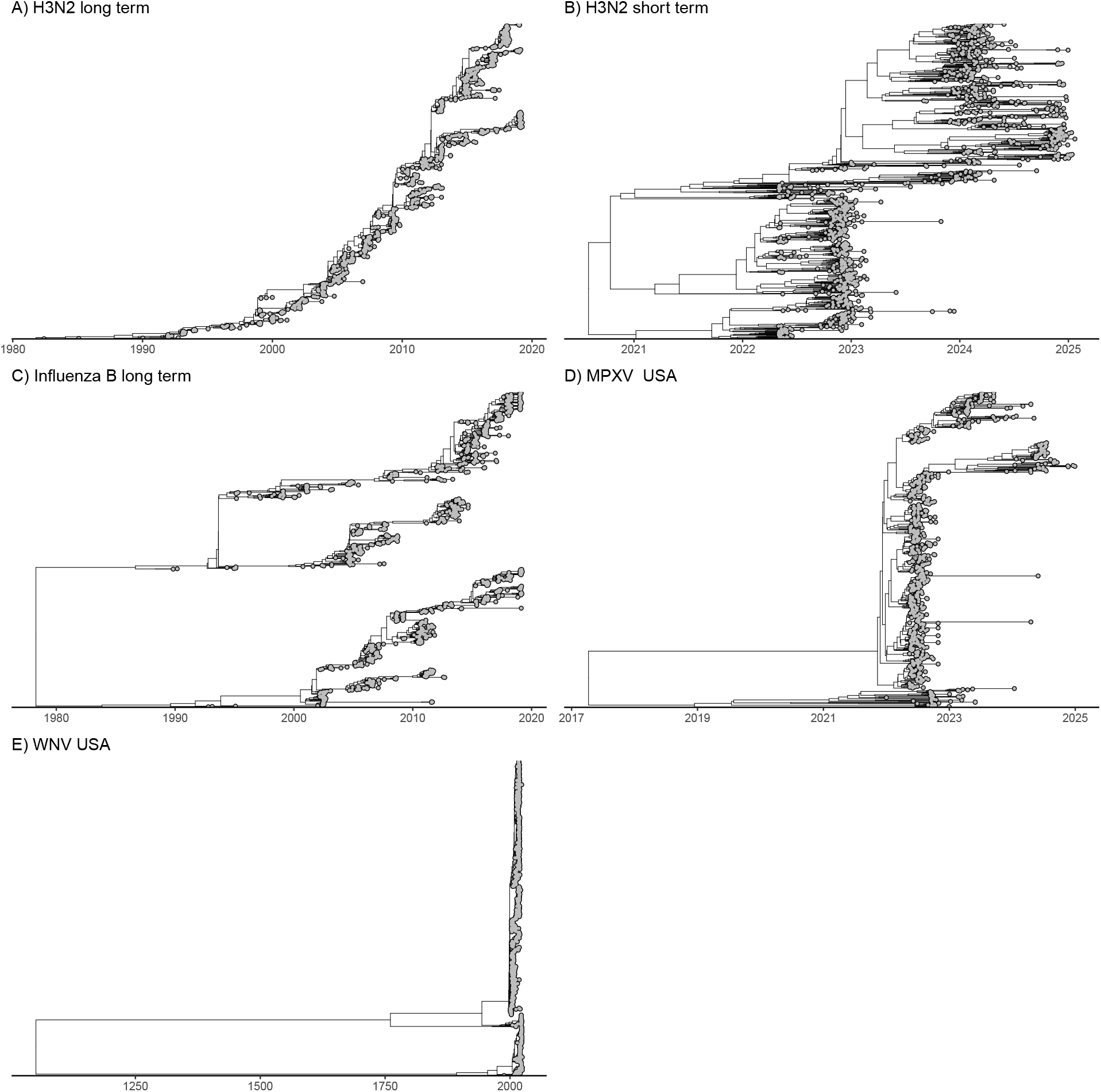
MCC trees off the human viral sequence data analyses. Here we show the maximum clade credibility tree for the convergence analyses of the real-world, viral datasets. The trees are intended to showcase both sampling intervals and tree shapes in the analyses.

**Figure S6:**
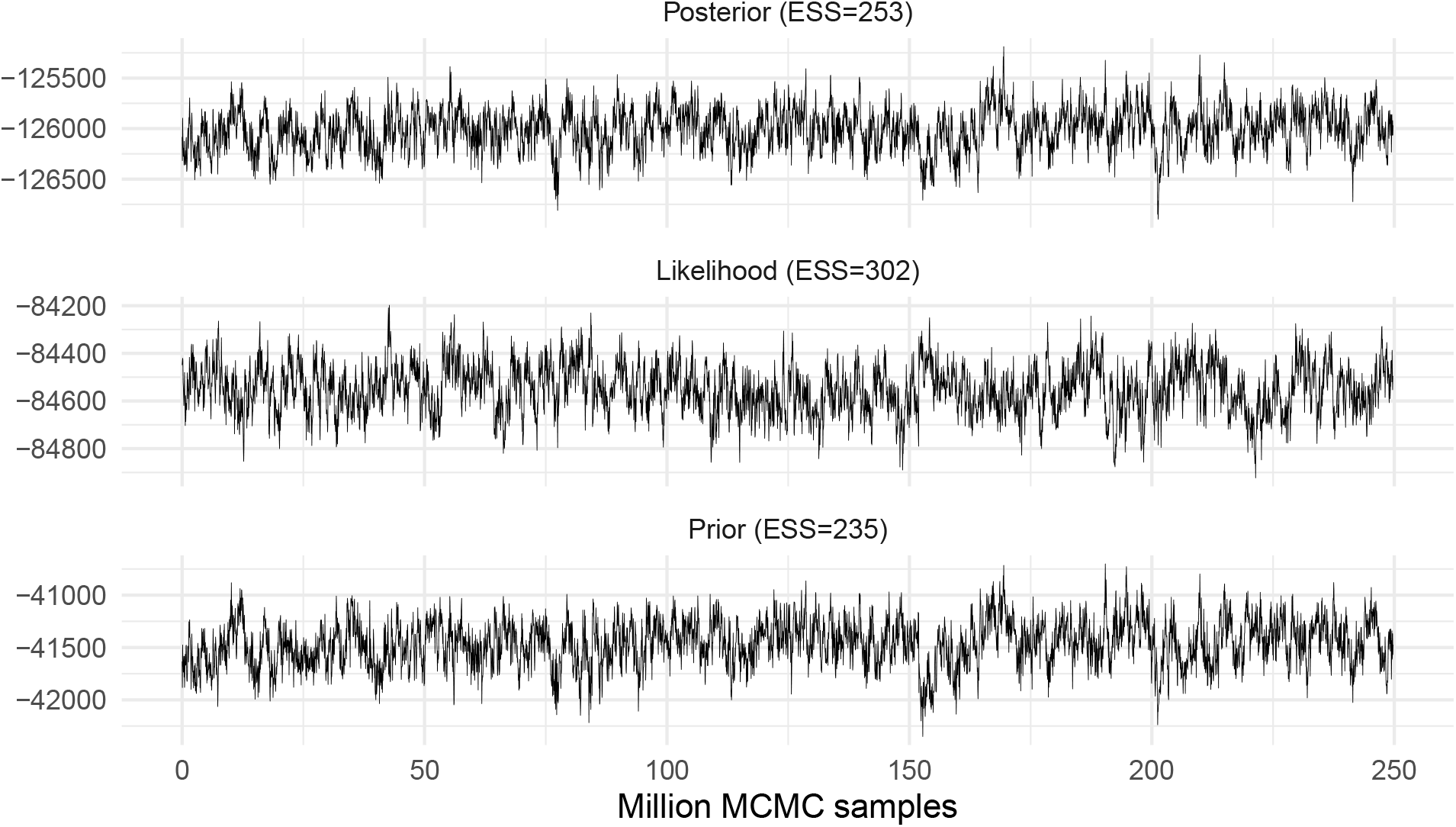
Traces for the Joint Bayesian demographic and discrete trait analysis with 10,000 sequences of H3N2. traces of posterior, likelihood, and prior combined from three analyses with indication of ESS.

**Figure S7:**
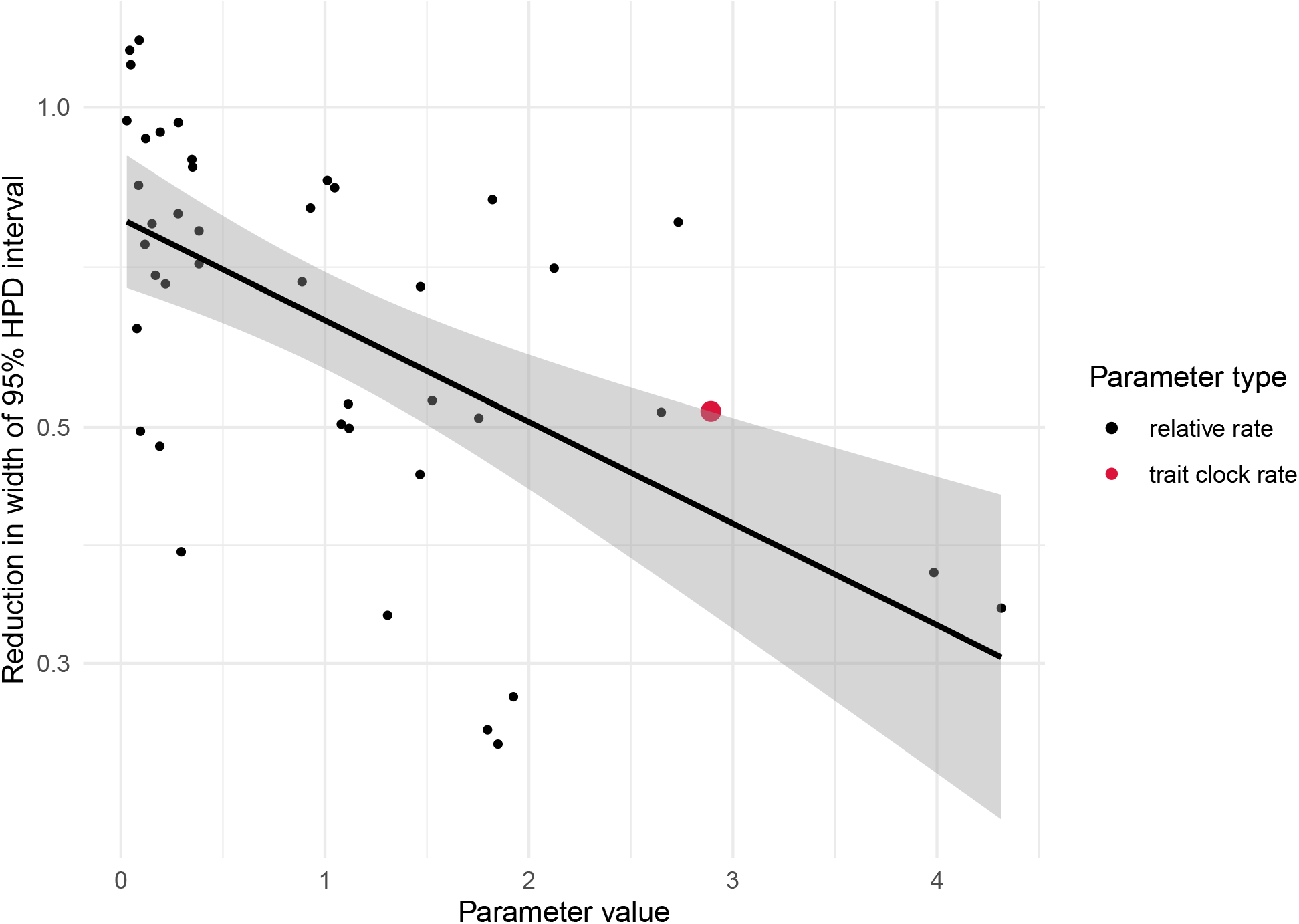
Improved precision of posterior estimates with increased data. Here, we compare the posterior interval widths for relative geographic rates and trait clock rates inferred using two datasets of different sizes (1,000 vs 10,000 sequences phylogeographic H3N2 analysis). Each point shows the log-scale reduction in the width of the 95% highest posterior density (HPD) interval (y-axis) as a function of the parameter’s posterior mean (x-axis), comparing the large dataset to the small one. A linear regression fit (black line) summarizes the overall trend. A higher value indicates greater precision achieved through increased data.

